# Five patterns of cell signaling pathways associated with cell behavior

**DOI:** 10.1101/2020.08.04.235986

**Authors:** Yuji Takeda, Kazuharu Kawano, Rui Ma, Shinichi Saitoh, Hironobu Asao

## Abstract

Cell signaling pathway is complex systems. Here, we present a concept for a new approach to analyze cell signaling pathway associated with cell behavior. In theoretically, cell behavior is recognized by energy and fluctuation. In this study, we measured phosphorylation level of signal transducers in a cell and fluctuation of the phosphorylation level in the cell population using flow cytometry. Flow cytometric data of mean fluorescence intensity (MFI) and coefficient variation (CV) were considered to the energy and the fluctuation, respectively. Topologically, the changes of MFI and CV were categorized into five patterns (we tentatively named as attractive, subsequent, passive, counter, and negative arbiter). In this study, we clarified the relationship between the cell behavior and the five patterns. Furthermore, combining the five patterns can define the signaling pathways, such as simple activated signal, oscillating signal, regulatory signal, robust signal, or homeostatic signal. These observations provide a proof of concept for general strategy to use the five patterns for connection between cell signaling pathway and cell behavior.

## Introduction

Life is a complex system that combines plasticity and robustness. How do we understand this system created by nature?

To find the answer, various signaling molecules have been identified and complex signaling mechanisms have been investigated (Krauss, 2014). In recent years, comprehensive (omics) analysis of various hierarchies, such as nucleic acids, proteins, or metabolites, has been performed, and many databases have been constructed. Recently, trans-omics, which combines the knowledge of signal transduction mechanisms, and database of omics analysis with systems biology, have been successful (Ohigashi *et al*, 2019; Yugi *et al*, 2016). However, these omics approaches presuppose that a cell population behaves uniformly.

Although the method of lysing (homogenization) a cell population is a powerful method for detecting the interaction or increase/decrease in a small number of molecules, a cell population is regarded as one large cell. As a result, homogenization explains the signal transduction pathway as a tight control model expressing robustness and regulation under a hypercomplex network.

According to reports on single-cell analysis, which have advanced in recent years, individual cell responses vary greatly (Raj and van Oudenaarden, 2008; Wimmers *et al*, 2018). Experimental fluctuations always occur in all experiments using biological materials. The fluctuation may be routinely considered as an error that is engendered by an undetectable initial difference or stochastic environmental difference even in a homogeneous and purified cell population. Are these fluctuations meaningless? To understand the complex system of life, we assumed that an epistemic tool for linkage between fluctuations and omics approaches should be prepared.

Previously, we measured the fluctuations of a cell population by staining a phosphorylated signal transduction molecule by flow cytometry (FCM) (Takeda *et al*, 2017). These previous observations allow us to speculate that the level of the fluctuations indicated emergent properties and robustness. In this study, we demonstrated that our approach provides a new concept for connection between cell signaling pathway and cell behavior.

## Results

### Theoretical model of cell behavior in space X

A cell receives many signals from the extracellular milieu and then responds to these signals. The behavior of a cell can be expressed in multi-dimensional space X, as shown by the X_1, 2… k_ axis (Fig 1A). These X axes are parameters that are defined by each experimental observation. A cell (small ball) freely moves in space X, and then the cell is attracted to a basin (W_1_ or W_2_) of attractors, indicating that the cell responds to some signals. Many attractors in space X perturbate the moving cell. The cell is trapped for a long time by strong attraction (dashed arrow of W_1_), indicating a robust cell response. However, weak attraction cannot trap a cell for a long time in the basin (dashed arrow of W_2_), indicating a susceptible cell response to other attractors. Therefore, weak attractors possess a large range of basins, and strong attractors possess a small basin range. The ranges of the basins are observed as fluctuations.

**Fig 1.**
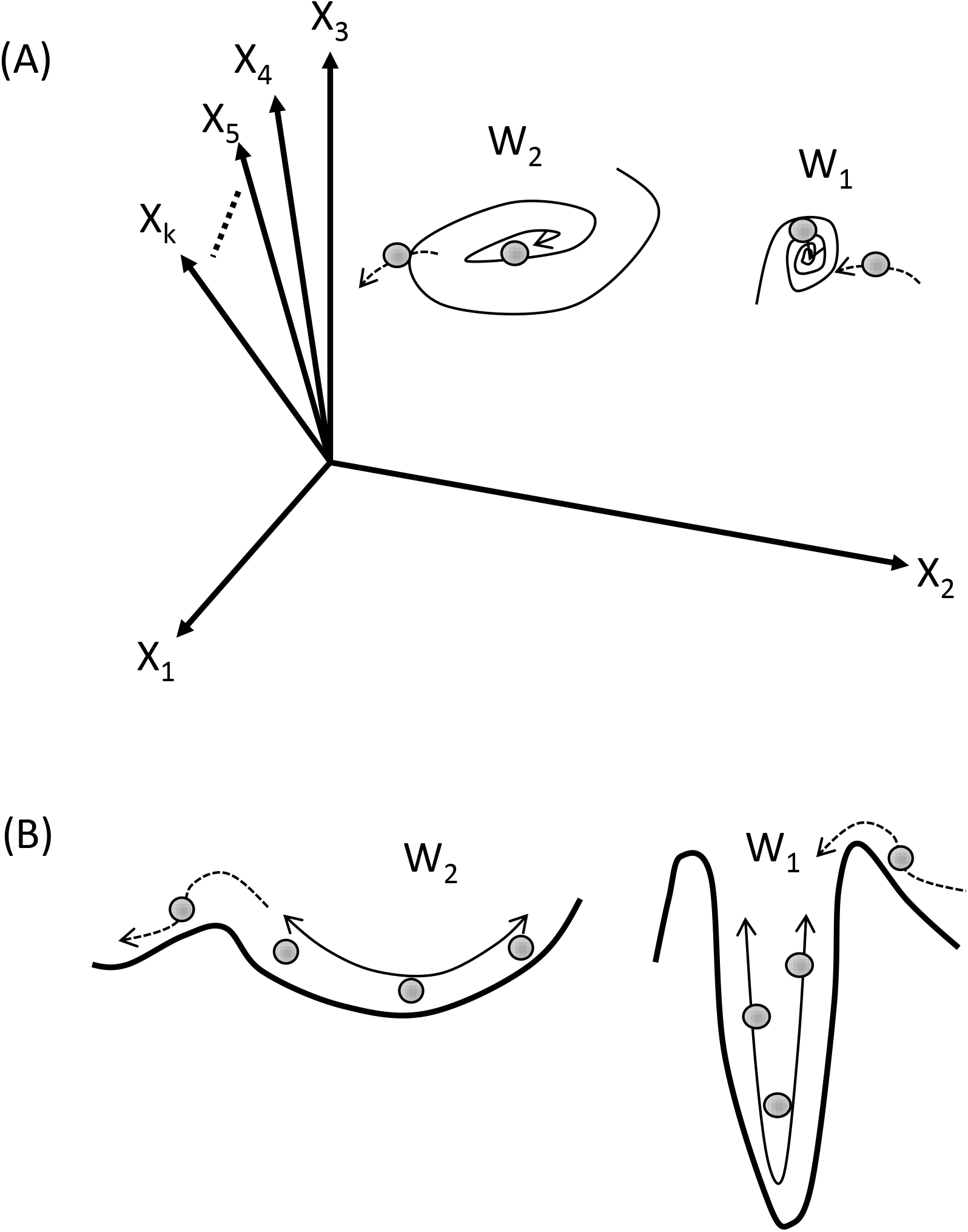
Schematic representation of relationship between cell behavior and attractor. Cells (gray balls) are freely moving in space X. When the cells respond to W_1_ or W_2_, cells are attracted to around W_1_ or W_2_. The trapping of cells into W_1_ or W_2_ are shown as whorled arrows. The basin of attraction is reflected in fluctuation of the moving cell. Transiting cells to another basin are indicated as dash arrows. The power of W_1_ is stronger than that of W_2_. (A) Schematic representation of cell behavior in the multi-parametric space of X. The attractive powers are indicated by white arrows. (B) Schematic representation of cell behavior on single parameter. The basins of attraction are presented as valleys and mountains. The attractive powers are presented as depth of valleys.

When the basins (W_1_ or W_2_) are observed from an appreciable axis, they present as valleys and mountains (Fig 1B). The power of attraction is shown as the depth of the valley. Removal to another valley requires a significant amount of energy, which is considered as the height of the mountain. This theoretical cell behavior has been proposed by previous research in biophysics (Furusawa and Kaneko, 2018). Furthermore, it is self-evident that the amount of energy is equivalent to the level of activated signaling pathways, such as ion influx, electron transfer, or protein phosphorylation, in life science.

### FCM analysis observations correspond to theoretical cell behavior

FCM is useful for the measurement of antigen expression per cell. When cells are stained with a fluorescence-conjugated antibody to the phosphorylated molecule of a signaling pathway, FCM analysis can define an activated level of the signaling pathway per cell. Furthermore, the sum of the data from each cell can be used to draw a histogram of the activated level in a cell population. Thus, the coefficient variation (CV) of the histogram indicates a fluctuation of each data point, corresponding to a basin of attractor. Furthermore, the mean fluorescence intensity (MFI) is the same as the mean value of the histogram, corresponding to a level of activated signaling pathway in a cell cluster.

It is generally assumed that histograms produce a total of five topologically distinct patterns compared to initial histograms (Fig 2A). The gray-filled histograms represent the initial status of a signaling pathway. During response to the signals, the histograms change, as shown in five patterns of open histograms (types 1–5) as follows: an increase in MFI and decrease in CV is type 1; an increase in both MFI and CV is type 2; both MFI and CV unchanged is type 3; a decrease in MFI and increase in CV is type 4; and a decrease in both MFI and CV is type 5.

**Fig 2.**
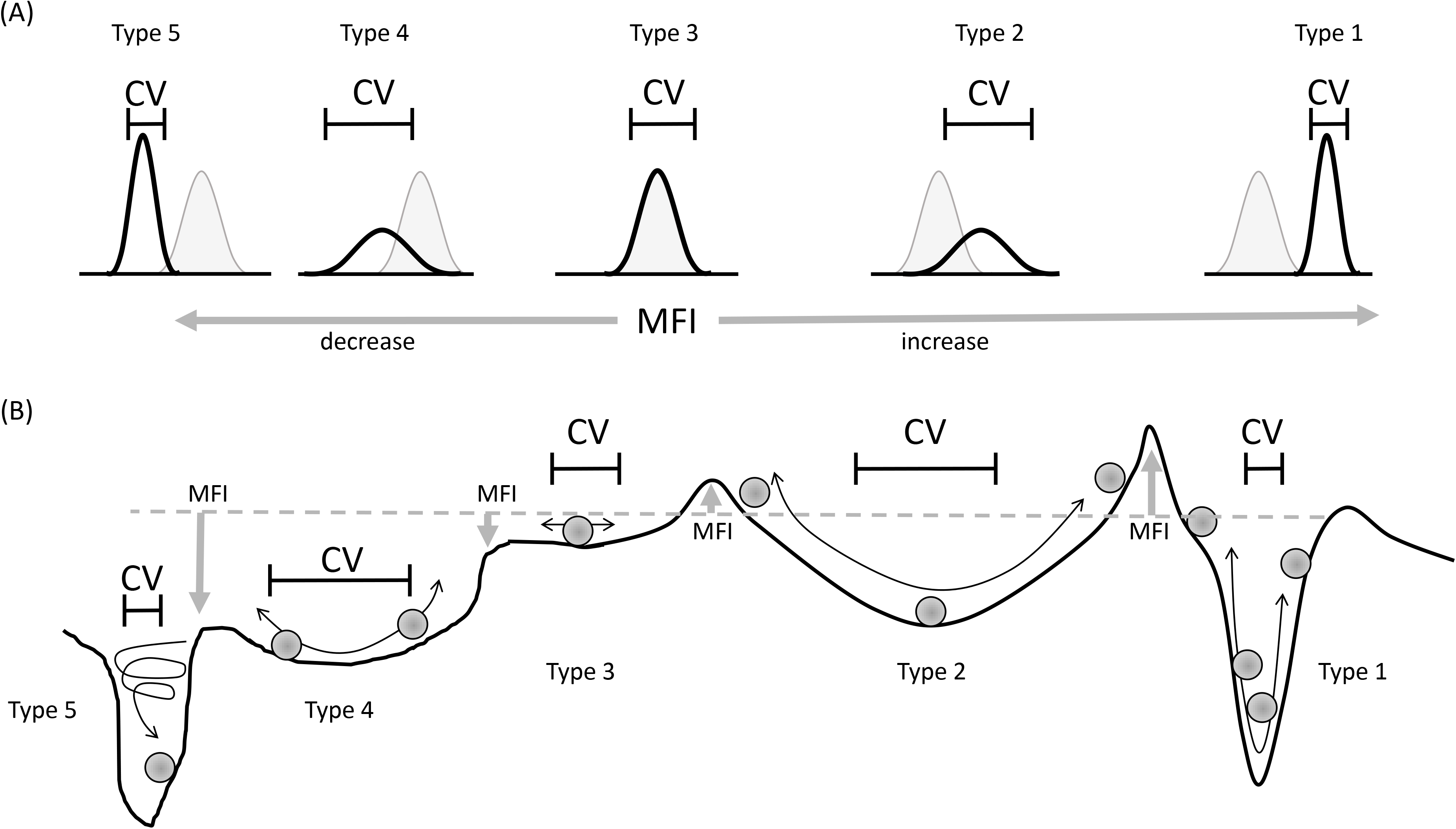
Schematic representation of the relationship between FCM analysis and theoretical cell behavior. (A) Five topological histogram patterns of FCM analysis. The gray-filled histograms and open histograms indicate the initial status and status after receiving signals, respectively. The histogram patterns after receiving signals are topologically categorized into five types (types 1–5) based on the increase or decrease in MFI (gray arrows) combined with an increase or decrease in CV (range of bars). (B) Relationship between FCM-analyzed parameters (MFI and CV) and theoretical cell behaviors. The theoretical cell behaviors are indicated in Figure 1B. A fluctuation in cells (basin) corresponds to a range of CV, and a significant amount of energy (height of mountain) is equivalent to the value of MFI. The changes in MFI and CV represent five types of cell statuses.

The relative change in the five histogram patterns is considered to be the five-cell status (Fig 2B). An increase in MFI compared to the initial level of MFI (dashed gray line) indicates that the cell receives a significant amount of energy to move into another basin. A decrease in MFI denotes the disappearance of a mountain, meaning that the decrease in energy is fusion with another basin. The power of attraction is shown as the depth of the valley, which is detected as CV. Therefore, type 1 is responsive to a robust attractor; type 2 is response to weak attractor; type 3 is in initial attractor; type 4 is response to weak and counteract attractor to initial attractor; and type 5 is level dropping to the ground state. In our study, we provisionally named the power of attraction as follows: type 1 is “attractive,” type 2 is “subsequent,” type 3 is “passive,” type 4 is “counter,” and type 5 is “negative arbiter” (Table 1).

**Table 1.** Categorization of cell status by FCM analysis.

| Categories | Change in MFI/CV | Provisional name of cell status |
| --- | --- | --- |
| Type 1 | MFI↑/ CV↓ | Attractive |
| Type 2 | MFI↑/ CV↑ | Subsequent |
| Type 3 | unchanged | Passive |
| Type 4 | MFI↓/ CV↑ | Counter |
| Type 5 | MFI↓/ CV↓ | Negative arbiter |

The categories correspond to Fig 2. “↑” or “↓” are indicated as increase or decrease, respectively.

**Table 2.** Type 3 (passive) can be separated from the other six statuses using change in MFI or CV and correlation between MFI and CV.

| Categories | Statistical<br>change in<br>MFI | Statistical<br>change in<br>CV | Correlation<br>between MFI<br>and CV | Speculative cell status |
| --- | --- | --- | --- | --- |
| **Type 1/2** | Increase | NS | NS | Intermediate between type 1 and 2<br>(mirror image of NF-κB in later phase) |
| **Type 2/4** | NS | Increase | NS | Perturbation by another basin<br>(p-tyrosine) |
| **Type 2/5** | NS | NS | Positive<br>correlation | Shuttling between type 2 and 5<br>(mirror image of homeostatic ERK1/2) |
| **Type 3** | NS | NS | NS | “Passive” as shown in Table 1 |
| **Type 1/4** | NS | NS | Negative<br>correlation | Shuttling between type 1 and 4<br>(Homeostatic ERK1/2) |
| **Type 1/5** | NS | Decrease | NS | Disappearance of perturbation by<br>another basin<br>(mirror image of p-tyrosine) |
| **Type 4/5** | Decrease | NS | NS | Intermediate between type 4 and 5<br>(NF-κB in later phase) |

NS, not significant. The categories correspond to Fig 11.

### Changes in CV shows that differences in cell behavior are dependent on stimulants

Previously, we measured several phosphorylated signal transducers by FCM (Takeda *et al*, 2014; Takeda *et al*, 2017). A previous report demonstrated that stimulants or inhibitors can induce the fluctuation of activated signals (change in CV) related to the level of phosphorylation (change in MFI) (Takeda *et al*, 2017). However, it was still unclear whether CV was dependent on the character (affinity or specificity) of the antibody and if it was regulated by the dose of stimulants.

To test effect of the character of the antibody and the dose of stimulants on CV, HL60 cells were stimulated with interferon (IFN)-α or granulocyte-colony stimulating factor (G-CSF), and then MFI and CV of phosphorylated STAT3 (pSTAT3) in the cells were measured by one kind of anti-phosphorylated STAT3 mAb. The representative flow cytometric data of HL60 cells are shown in Fig EV 1A‒C. IFN-α increased pSTAT3 MFI with a peak at 30 min (Fig 3A). Similarly, G-CSF increased pSTAT3 MFI with a peak at 10–30 min (Fig 3A). The same anti-pSTAT3 mAb was used for these experiments. However, G-CSF significantly increased pSTAT3 CV with a peak at 10 min, while IFN-α did not increase pSTAT3 CV during the incubation time (Fig 3B). Therefore, these results show that the changing pattern of CV during cell activation is dependent on the stimulants but not on the character of the antibody.

**Fig 3.**
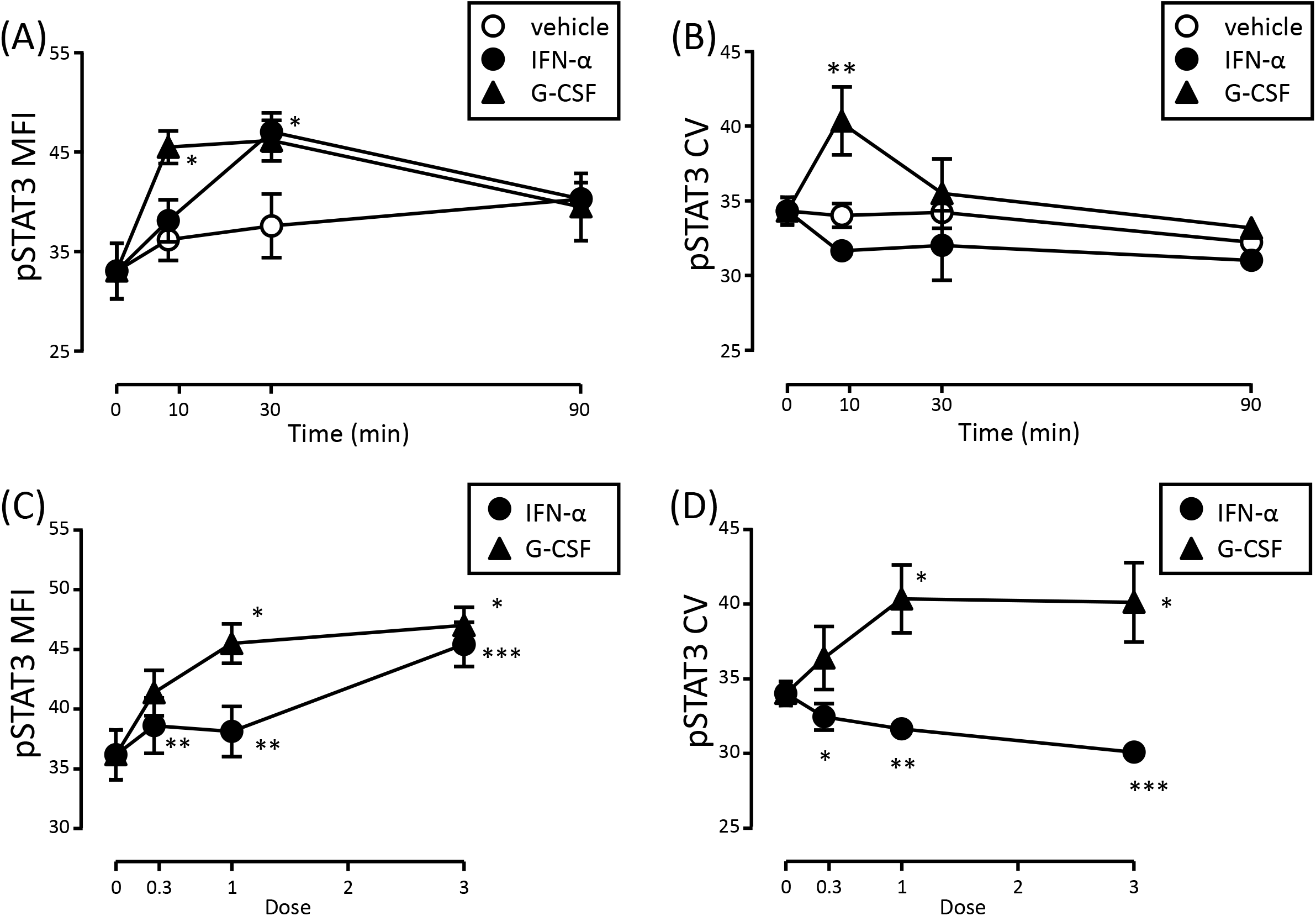
Change in MFI and CV after stimulation with IFN-α or G-CSF. HL60 cells were stimulated with vehicle (open circle), IFN-α (closed circle), or G-CSF (closed triangle). The cells were stimulated with IFN-α (100 U/mL) or G-CSF (1 nM) for 0, 10, 30, and 90 min (A and B). The cells were stimulated with the various doses of IFN-α (0.3, 33 U/mL; 1, 100 U/mL; 3, 300 U/mL) or G-CSF (0.3, 0.33 nM; 1, 1 nM; 3, 3 nM) for 10 min (C and D). After the stimulations, the cells were fixed/permeabilized and stained with Alexa488-conjugated anti-phosphorylated STAT3 mAb. The cells were measured by FCM (ec800), and then MFI (A and C) and CV (B and D) of the histograms were analyzed. Data are represented as mean ± SE from three independent experiments. Statistical analysis used two-way ANOVA with post-hoc test using Bonferroni’s correction for comparison to vehicle in (A) and (B) or one-way ANOVA with post-hoc test using Bonferroni’s correction for comparison to dose of 0 in (C) and (D). \**p* < 0.05; \*\**p* < 0.01; \*\*\**p* < 0.001.

Next, we investigated whether different cytokine concentrations cause different changes in MFI and CV. Cells were stimulated with several doses of IFN-α (33, 100, or 300 U/mL) or G-CSF (0.33, 1, or 3 nM) for 10 min. Stimulation with both IFN-α and G-CSF significantly increased pSTAT3 MFI in a dose-dependent manner (Fig 3C). However, IFN-α and G-CSF induced opposite changes in the pSTAT3 CV, with IFN-α decreasing pSTAT3 CV but G-CSF increasing pSTAT3 CV (Fig 3D). These results indicated that the direction of CV change was dependent on the type of stimuli but not on the dose of stimulants even in the same cell line.

### Signaling pathway containing transient activation can be distinguished by correlations between MFI and CV

Activation of the signaling pathway is generally transient and simple. Although statistically significant changes in both MFI and CV are easily detected at the peak of activation, it is unclear that the significant changes in both MFI and CV can be detected before or after the peak of activation. In logically, when a cell population changes to a type 1 histogram from the initial histogram, the increase in MFI is accompanied with a decrease in CV, and when a cell population changes to a type 2 histogram from the initial histogram, the increase in MFI is accompanied by an increase in CV. Consequently, a negative correlation between MFI and CV will be detected in type 1 (attractive). While a positive correlation between MFI and CV will be detected in type 2 (subsequent). Therefore, it is possible that transient and simple activation may be detected by the correlation between MFI and CV independent of observation time.

To test these possibilities, the correlation between MFI and CV was analyzed. pSTAT3 MFI in cells stimulated with IFN-α or G-CSF increased transiently (Fig 3). IFN-α slightly decreased CV and showed a negative correlation between pSTAT3 MFI and CV (Fig 4A). G-CSF showed a transient increase in CV and showed a positive correlation between pSTAT3 MFI and CV (Fig 4B). These analyses provided evidence that the signaling pathway containing transient and simple activation can be distinguished by the correlation between MFI and CV. Thus, at any time point or during the sum of time points, type 1 (attractive) exhibited a negative correlation and type 2 (subsequent) exhibited a positive correlation.

**Fig 4.**
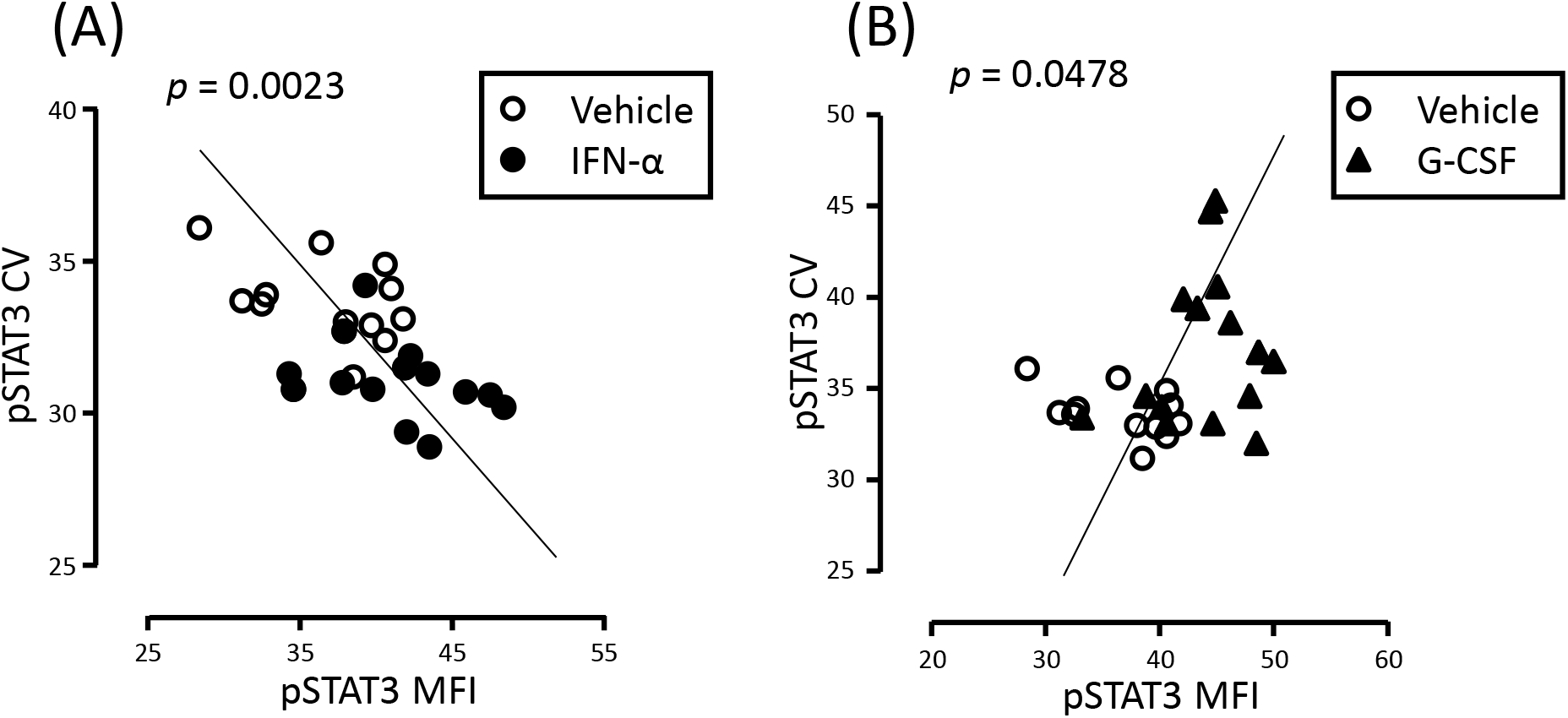
Correlations between MFI and CV during stimulation with IFN-α or G-CSF. HL60 cells were stimulated with vehicle (open circle; A and B), IFN-α (closed circle; A), or G-CSF (closed triangle; B), similar to the conditions shown in Fig 3. The cells were measured by FCM (ec800), and then pSTAT3 MFI (horizontal axis) and pSTAT3 CV (vertical axis) were analyzed. The data consisted of three independent experiments. The statistical significances of correlation were calculated with Pearson’s correlation coefficient (*p* value is indicated in each figure).

Because types 4 and 5 are mirror images of types 2 and 1, the positive or negative correlations can also be applied to transient deactivation (i.e., temporal decrease in MFI) to type 4 (counter) or type 5 (negative arbiter), respectively.

### Signaling pathway consisting of oscillating activation can be recognized as the persistence of type 2 (subsequent) without the positive correlation between MFI and CV

Activation of NF-κB is oscillatory-controlled to regulate its biological functions (Paszek *et al*, 2010). How can this oscillating activation signaling pathway be recognized by FCM?

The oscillation pattern is known that the level of NF-κB activation in the early phase (*t*_*1*_) is higher than that in the later phase (*t*_*2*_) (Fig 5A). Thus, the oscillation pattern indicates that the cell moving from the initial basin (W_0_) to another basin (W_1_) will occur in the early phase (*t*_*1*_) but not in the later phase (*t*_*2*_), as shown in conceptual schema (Fig 5B). If the oscillation is highly synchronized in a cell population, the oscillation signal should be simply detected as oscillatory changes in MFI without changes in CV. However, the oscillatory changes are not generally synchronized in a cell population. Therefore, the amplitudes should be accompanied with an increase in CV (Fig. 5B), indicating that the persistence of oscillation should be recognized as an increase in CV compared to the initial status of CV (*t*_*0*_ vs *t*_*1*_ or *t*_*2*_; Fig 5B). Furthermore, the amplitudes in the later phase is smaller than the amplitudes in early phase (Fig. 5A), indicating that MFI in the later phase should be decrease, compared to MFI in the early phase (*t*_*1*_ vs *t*_*2*_; Fig 5B). In summary, the oscillation signal will be detected as the persistence of an increase in both MFI and CV (type 2) containing a decrease in MFI in later phase.

**Fig 5.**
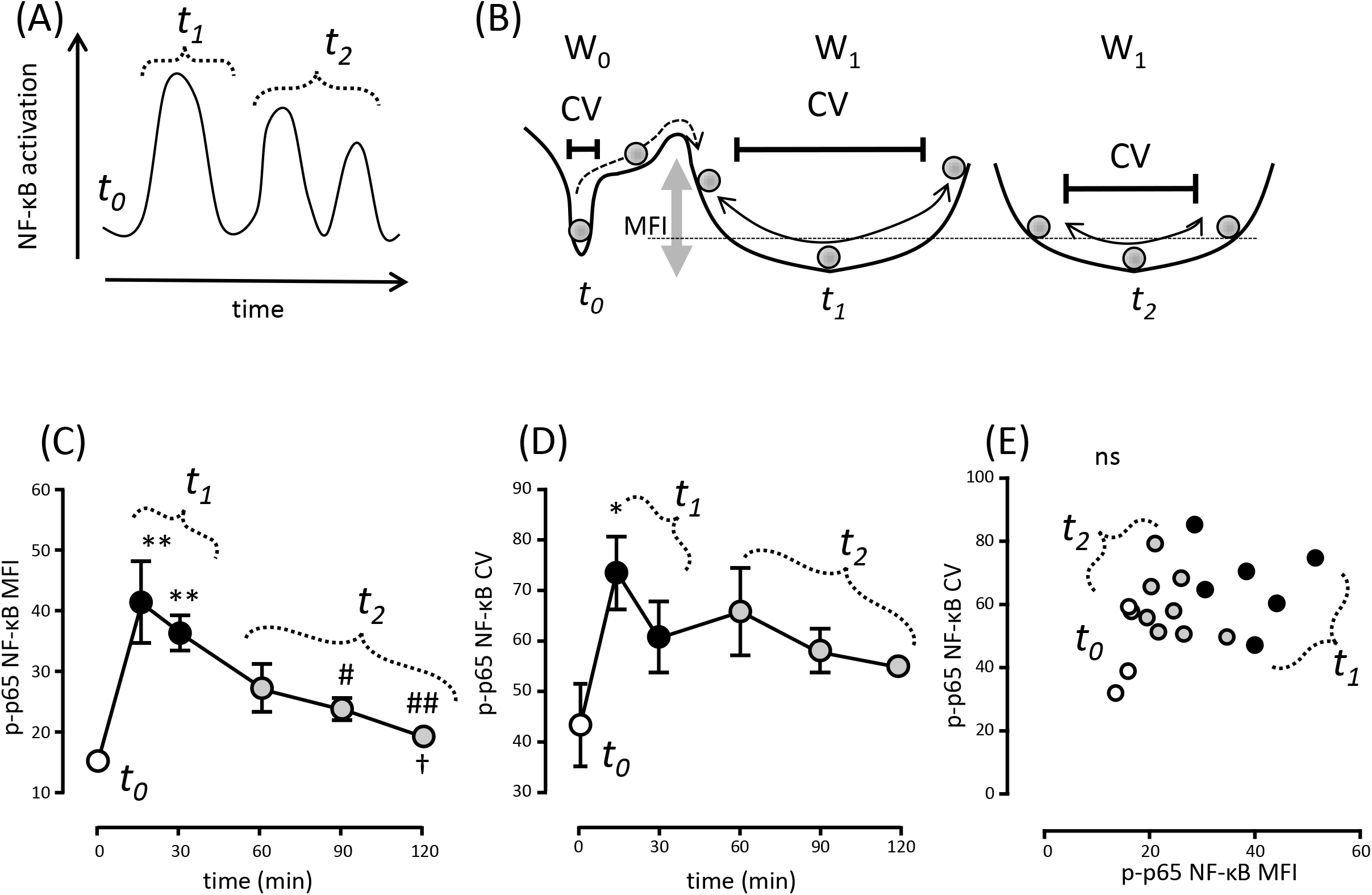
Analysis of oscillatory signal of NF-κB using FCM. (A) Schematic representation of oscillatory NF-κB activation according to a previous report. After the stimulation, NF-κB was quickly activated in the early phase (*t*_*1*_) from the initial status (*t*_*0*_). The activation gradually decreased in the later phase (*t*_*2*_). (B) Schematic representation of the relationship between oscillatory cell behavior and FCM analysis (MFI and CV). The transition of the phosphorylated cell into an oscillatory basin (W_1_, *t*_*1*_) from the initial basin (W_0_, *t*_*0*_) is indicated by dashed arrows in the early phase. The amplitudes in the oscillatory basin (W_1_) decreased depending on the level of phosphorylation in the later phase (*t*_*2*_). (C and D) Changes in phosphorylated p65 NF-κB (p-p65 NF-κB) in activated U937 cells. The cells (2 × 10^5^ cells/mL) were stimulated with PMA (100 ng/mL) for 15, 30, 60, 90, and 120 min. After stimulation, the cells were fixed/permeabilized and stained with rabbit anti-p-p65 NF-κB mAb and Alexa488-conjugated goat anti-rabbit polyclonal Ab. The cells were measured by FCM (ec800), and the changes in p-p65 NF-κB MFI (C) and p-p65 NF-κB CV (D) were analyzed. Statistical significance was calculated by one-way ANOVA with post-hoc by Tukey’s test (n = 3), \**p* < 0.05, \*\**p* < 0.01 (compared to 0 min); ^#^*p* < 0.05, ^##^*p* < 0.01 compared to 15 min; ^†^*p* < 0.05 compared to 30 min). There was no significant difference among the other pairs (no indication). The statistical significance of correlations was calculated with Pearson’s correlation coefficient (not significant, ns).

To verify this hypothesis, the levels of phosphorylated p65 NF-κB (p-p65 NF-κB) MFI and p-p65 NF-κB CV were measured in U937 cells stimulated by phorbol 12-myristate 13-acetate (PMA). The representative flow cytometric data of U937 cells are shown in Fig EV 1D‒F. The increase in p-p65 NF-κB MFI stimulated by PMA was accompanied with an increase in p-p65 NF-κB CV in the early phase (*t*_*0*_ compared to *t*_*1*_; Fig 5C and D). After the early phase, the level of p-p65 NF-κB MFI decreased significantly depending on the course of time in the later phase (*t*_*1*_ compared to *t*_*2*_; Fig 5C). Similarly, p-p65 NF-κB CV decreased in the later phase but not significantly (*t*_*1*_ compared to *t*_*2*_; Fig 5D). These changes in MFI and CV were categorized as follows: the change between *t*_*0*_ and *t*_*1*_ was an increase in both MFI and CV, which was typical type 2 (subsequent); the change between *t*_*1*_ and *t*_*2*_ was a decrease in MFI; and the change between *t*_*0*_ and *t*_*2*_ was an increase in both MFI and CV, which was type 2 (subsequent). Overall, the oscillation signal was recognized as an increase in both MFI and CV, which was type 2 (subsequent). In the whole observed term from *t*_*0*_ to *t*_*2*_, there was no correlation between MFI and CV (Fig 5E) because type 2 and type 4/5 (intermediate between counter and negative arbiter) were mixed in the term. These results indicated that the categorization using FCM can recognize the oscillating signaling pathway as “type 2 and no correlation”.

### Homeostatic ERK1/2 pathway is a special pattern of type 3 (passive) with a constant negative correlation between MFI and CV

Type 3 (passive) belongs to a category in which there is no significant difference between the stimulatory condition and initial state (Fig 2). Previously, we reported that a negative correlation between phosphorylated ERK1/2 (pERK1/2) MFI and pERK1/2 CV was detected in Jurkat cells without significant changes in both MFI and CV (Takeda *et al*, 2017).

In this study, we measured pERK1/2 using human T cells and neutrophils in whole peripheral blood stimulated with various cytokines, such as IFN-α, interleukin (IL)-21, and G-CSF alone or in combination, to confirm the negative correlation. Although IFN-α and IL-21 primarily activate STAT3 pathways, these cytokines simultaneously activate the ERK pathway in various types of cells (Juliana *et al*, 2012; Zhao *et al*, 2015). G-CSF also primarily activates STAT3 and concomitantly activates the ERK pathway in myeloid cells (Kamezaki *et al*, 2005). Whole blood was stimulated with these cytokines, and then pERK1/2 levels were analyzed in CD3^+^ cells (T cells) or GPI-80^+^ cells (neutrophils). The representative flow cytometric data of T cells and neutrophils in peripheral blood are shown in Fig EV 2. The various stimulations and combinations did not significantly change both MFI and CV of pERK1/2 in T cells (Fig 6A and B) and neutrophils (Fig 6C and D). These results indicate that the category of ERK1/2 pathway in both T cells and neutrophils is type 3 (passive) under all tested stimulations. Interestingly, these data showed significantly negative correlations between pERK1/2 MFI and pERK1/2 CV (Fig 6E and F). The negative correlation without a significant increase in MFI indicated the existence of persistently controlled signals without an increase in the moving energy. It is well-known that the ERK pathway, which plays a crucial role in cell survival, induces asynchronous pulses of varying frequencies and durations (Albeck *et al*, 2013; Chambard *et al*, 2007). Thus, the negative correlation without increase in MFI may indicate cell survival signals, i.e., “homeostatic” ERK pathway.

**Fig 6.**
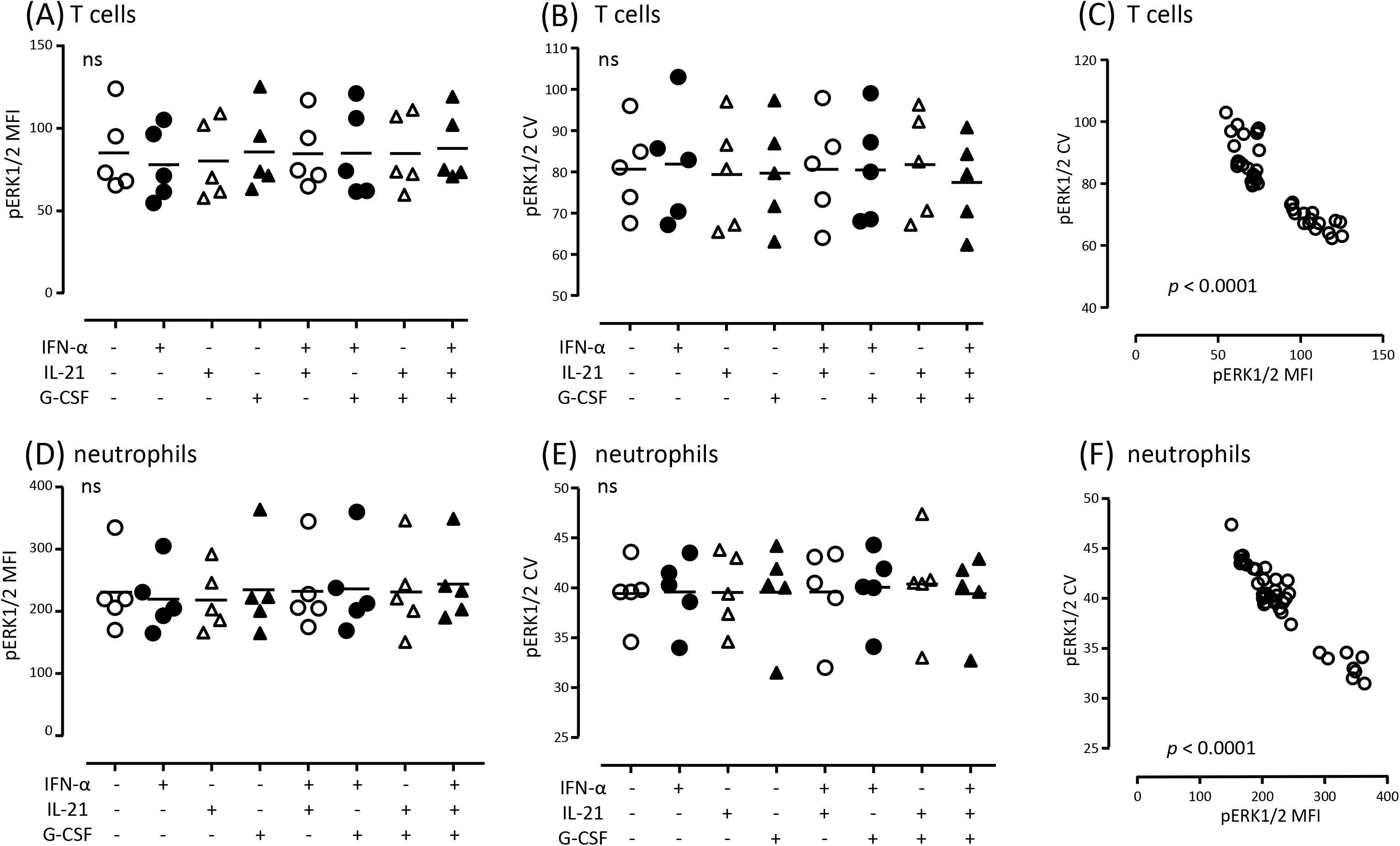
Analysis of homeostatic ERK pathway using FCM in T cells and neutrophils. Human peripheral blood cells were stimulated with IFN-α (100 U/mL), IL-21 (1 nM), G-CSF (1 nM), or a combination of these cytokines for 30 min. After the stimulations, the cells were fixed/permeabilized and stained with Pacific Blue-conjugated anti-human CD3 mAb, PE-conjugated anti-GPI-80 mAb, and Alexa647-conjugated anti-phosphorylated ERK1/2 mAb. The cells were measured by FCM (FACSCanto II). CD3^+^ cells were gated for T cell analysis (A and B), and GPI-80^+^ cells were gated for neutrophil analysis (D and E). MFI and CV of phosphorylated ERK1/2 (pERK1/2) in each gated cell population were analyzed. The correlations between MFI and CV on T cells and neutrophils are shown in (C) and (F), respectively. The data consisted of more than four independent experiments, and the statistical significances of correlation were calculated with Pearson’s correlation coefficient (*p* value is indicated in each figure).

To verify that type 3 (passive) with negative correlation is the homeostatic ERK pathway, we compared the emergent response via the ERK pathway, such as inflammatory responses induced by lipopolysaccharide (LPS) (Guha and Mackman, 2001; Nick *et al*, 1996). Whole blood was stimulated with LPS, and then pERK1/2 was analyzed in CD14^+^ monocytes. The representative flow cytometric data of monocytes in peripheral blood are shown in Fig EV 3. Both pERK1/2 MFI and pERK1/2 CV increased significantly by LPS stimulation, i.e., type 2 (subsequent) (Fig 7A). Furthermore, the correlation between pERK1/2 MFI and pERK1/2 CV was not detected (Fig 7B), suggesting that the “subsequent and no correlation” was oscillated activation similar to NF-κB pathway in U937 cells stimulated with PMA (Fig 5). Indeed, the ERK pathway is known as an oscillating signal (Shankaran and Wiley, 2010). These results indicate that the ERK pathway shows two different types of category, oscillating type 2 and homeostatic type 3, depending on the stimulant and cells.

**Fig 7.**
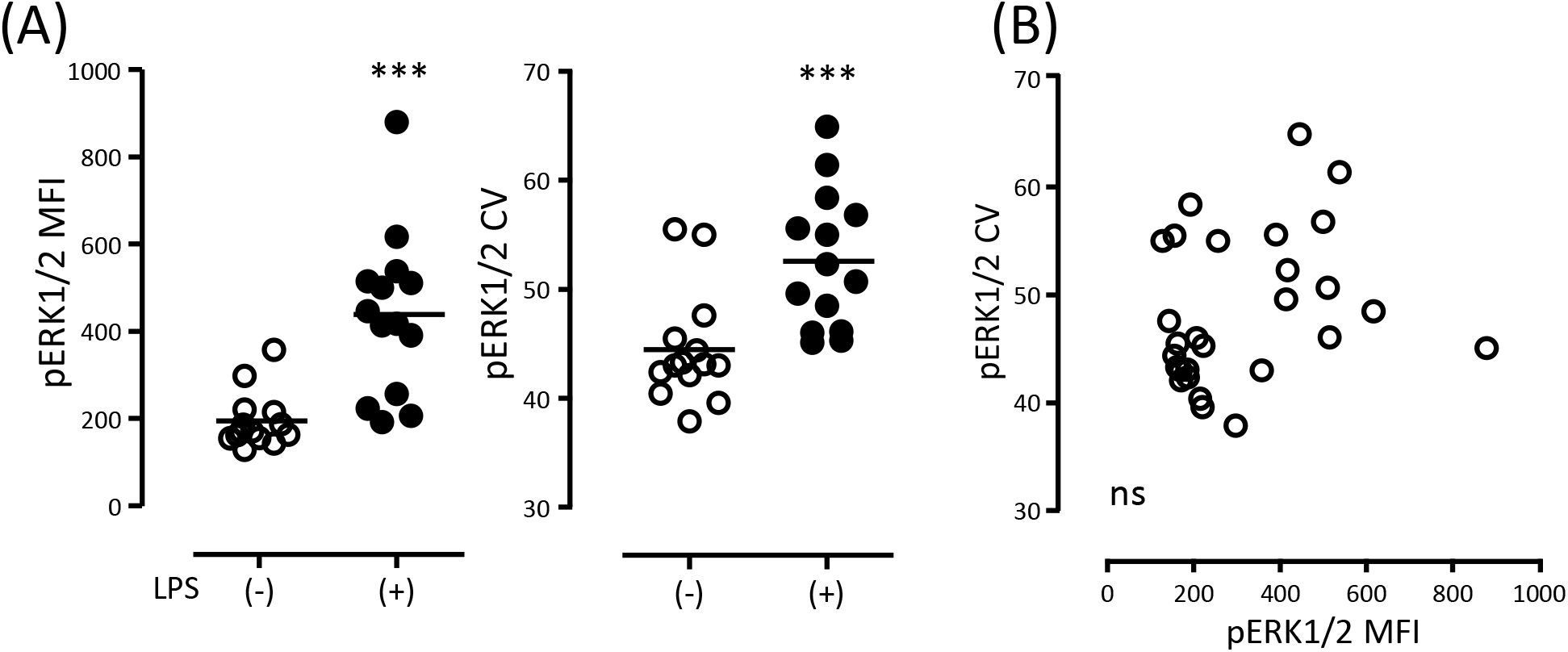
Analysis of emergent ERK pathway in monocytes using FCM. Human peripheral blood cells were stimulated with (+) or without (−) LPS (1 μg/mL) for 30 min. After the stimulation, the cells were fixed/permeabilized and stained with FITC-conjugated anti-human CD14 mAb and Alexa647-conjugated anti-phosphorylated ERK1/2 (pERK1/2) mAb. (A) MFI and CV of phosphorylated ERK1/2 (pERK1/2) in gated CD14^+^ monocytes were measured using FCM (FACSCanto II). Data were obtained from four independent experiments, and the statistical significance was calculated by Wilcoxon signed-ranks test, \*\*\**p* < 0.001 compared to LPS (−). (B) The correlations between MFI and CV are indicated. Data were obtained from four independent experiments, and the statistical significance was calculated by Pearson’s correlation coefficient (not significant, ns).

### Detection of regulatory and robust signaling pathways

The network of signaling pathways consists of many signaling modules. These signaling modules act coordinately or independently. The interaction of these modules is considered to induce variety and plasticity responses (Krauss, 2014). Can the categorization using FCM be useful to detect the coordinate modules (regulatory signaling pathways) or independent modules (robustness signaling pathways)?

Theoretically, robustness is a small basin and perturbated fluctuation is a large basin (Fig 2), indicating that robustness or regulatory pathways will be found as type 1 or type 2, respectively. Previously, it has been demonstrated that the phosphatidylinositol 3-kinase-Akt pathway limits LPS activation of signaling pathways (i.e., MAPK/ERK pathway) in human monocytic cells (Guha and Mackman, 2002). Therefore, we speculated that the Akt pathway is detected as a robust signal pathway, and that the MAPK/ERK pathway, which is regulated by the Akt pathway, is recognized as a perturbated regulatory signal pathway.

To verify the correspondence between change in CV and these types of signaling pathways, phosphorylation of Akt and p38 MAPK in monocytes were analyzed using FCM (Fig 8). After LPS stimulation, changes in phosphorylated p38 MAPK (p-p38MAPK) were recognized as type 2 (subsequent), which is a category with an increase in both MFI and CV, similar to that observed for pERK1/2 (Fig 7A and Fig 8A). Phosphorylated Akt (pAkt) was detected as type 1 (attractive), which is a category with an increase in MFI and decrease in CV (Fig 8B). These results suggested that our speculation was consistent with previous observations of LPS signaling modules.

**Fig 8.**
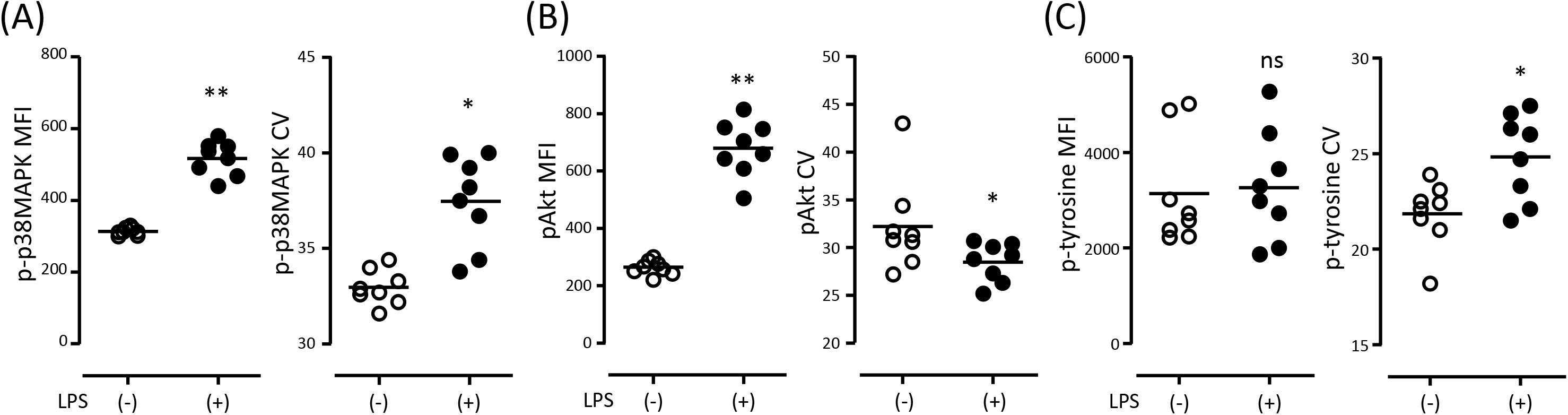
Analysis of regulatory and robust signaling pathway in monocytes using FCM. Human peripheral blood cells were stimulated with (+) or without (−) LPS (1 μg/mL) for 30 min. After the stimulation, the cells were fixed/permeabilized and stained with PE-conjugated anti-CD14 mAb, Alexa488-conjugated anti-phosphorylated p38 MAPK (p-p38MAPK) mAb, V450-conjugates anti-phosphorylated Akt (pAkt) mAb, and Alexa647-conjugated anti-phosphorylated tyrosine (p-tyrosine) mAb. MFI and CV of p-p38MAPK (A), pAKT (B), or p-tyrosine (C) in gated CD14^+^ monocytes were measured using FCM (FACSCanto II). Data were obtained from four independent experiments, and the statistical significance was calculated by Wilcoxon signed-ranks test, \**p* < 0.05; \*\**p* < 0.01; \*\*\**p* < 0.001 compared to LPS (−).

Furthermore, we analyzed total phosphorylated tyrosine (p-tyrosine) in monocytes. Conceptually, an increase in the module interaction can be imaged as the appearance of several basins around the initial basin or a decrease in the height of the mountain (fusion of basins). Many signal modules have p-tyrosine, and the modules must increase the interaction with each other during the emergent response, such as stimulation with LPS. Therefore, we examined whether p-tyrosine CV increased after the interaction of signal modules induced by LPS in monocytes.

The total level of p-tyrosine (p-tyrosine MFI) did not alter significantly (Fig 8C). This result is very reasonable because p-tyrosine MFI presents the sum of many kinds of signal modules containing phosphorylated and dephosphorylated tyrosine. However, p-tyrosine CV increased significantly (Fig 8C). These results demonstrated that the increase in CV is adaptable for the increase in signal module interaction.

### Identification of signaling pathways related to cell differentiation

What can the categorization using FCM be used for? The categorization may be used to predict a relatively robust signaling pathway among various signaling pathways. Such a robust signal should lead to downstream responses in the analyzed cell population. To assess the application of the categorization, we investigated the relationship between T cell differentiation and signaling pathways using FCM.

Ovalbumin (OVA) can stimulate CD4^+^ T cells in OT-II mice, which is a well-established system to examine T cell response via TCR signals (Barnden *et al*, 1998; Leung *et al*, 2013). In this study, we did not use attracting cytokines, such as IL-4 and IFN-γ, and analyzed the differentiated CD4^+^ T cells induced by OVA *in vitro*. The differentiation of T cells occurred randomly (attractive signal was statistically determined), and the differentiated Th1 cell subset was detected by the transcription factor T-bet, which is a master regulator of Th1 cells (Oestreich and Weinmann, 2012).

The T-bet^+^ cell subset (Th1) in CD4^+^ T cells was gated and compared to the T-bet^−^GATA3^−^RORγt^−^Bcl6^−^ cell subset in CD4^+^ T cells (Th0) (Fig 9). The representative flow cytometric data of monocytes in peripheral blood are shown in Fig EV 4. The increase in pSTAT3 MFI and decrease in pSTAT3 CV in Th1 were significant compared to that in Th0 (Fig 9A). Additionally, a negative correlation between pSTAT3 CV and pSTAT3 MFI was also observed. These results suggested that the STAT3 pathway was type 1 (attractive) for differentiation into Th1 from Th0. Similarly, the pERK pathway was also recognized as type 1 (attractive) (Fig 9B).

**Fig 9.**
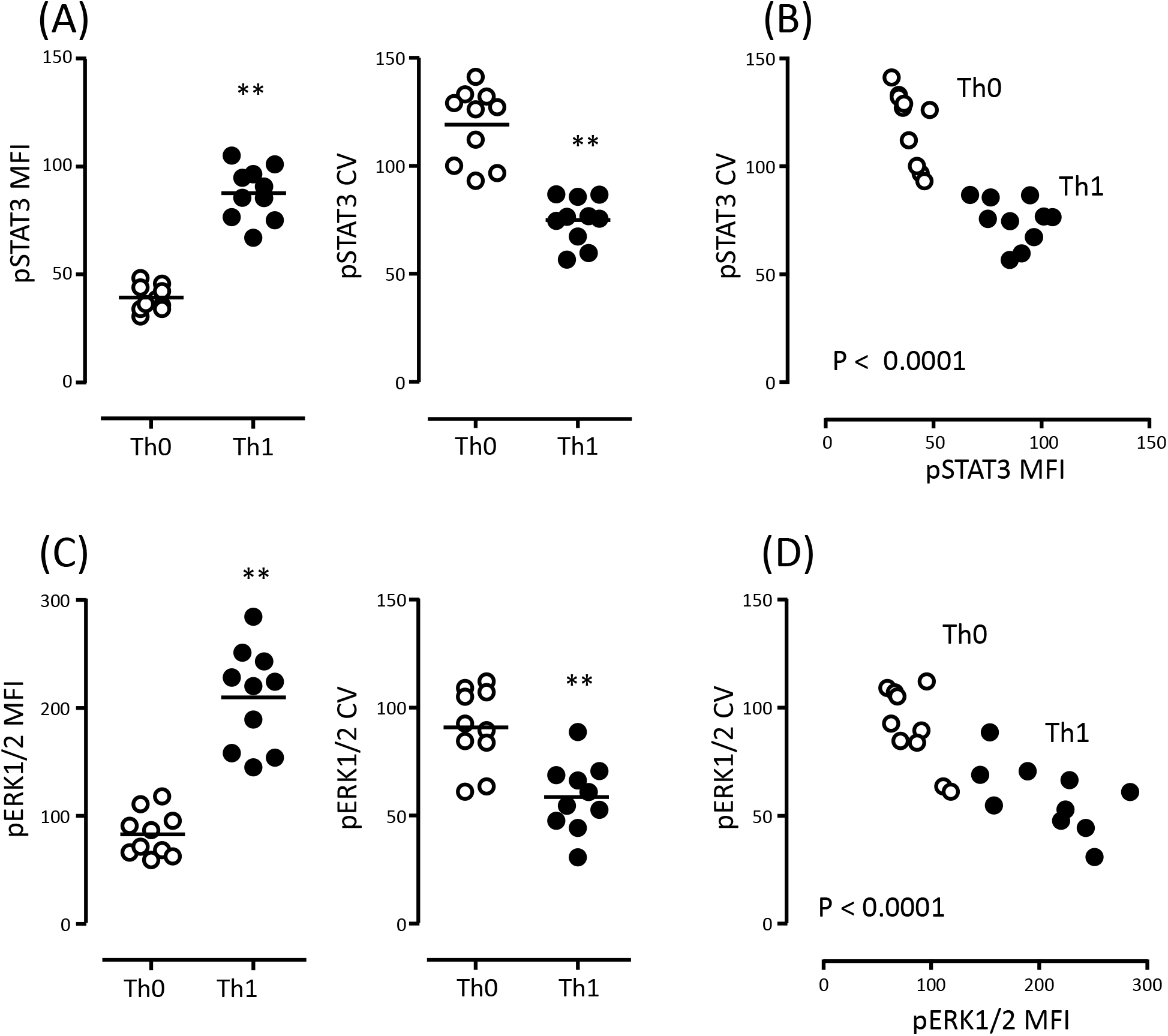
Analysis of STAT3 and ERK1/2 signaling pathways in Th0 and Th1 cells using FCM. Splenocytes were stimulated with OVA for 7 days as described in “Methods”. After the stimulations, the cells were fixed/permeabilized and stained with mAbs (V450-conjugated anti-mouse CD3 mAb, PE-Cy7-conjugated anti-mouse CD4 mAb, Alexa488-conjugated pSTAT3 mAb, Alexa647-conjugated pERK1/2 mAb, and PE-conjugated anti-T-bet mAb or a mixture of PE-conjugated anti-T-bet mAb, anti-GATA3 mAb, anti-RORγt mAb, and anti-Bcl-6 mAb). Cells were analyzed by FCM (FACSCanto II). Th0 cell subset was gated as T-bet^−^GATA3^−^RORγt^−^Bcl-6^−^ in CD3^+^CD4^+^ cells (open circle), and Th1 cell subset was gated as T-bet^+^ in CD3^+^CD4^+^ cells (closed circle). MFI and CV of pSTAT3 (A) or pERK1/2 (C) and the correlation between MFI and CV of pSTAT3 (B) or pERK1/2 (D) are presented. Data were obtained from more than four independent experiments, and the statistical significance was calculated by Wilcoxon signed-ranks test. \*\**p* < 0.01. The statistical significances of correlation were calculated with Pearson’s correlation coefficient (*p* value is indicated in each figure).

Which of the signaling pathways contributed to T cell differentiation? To compare the contribution, increase in the percentage (increase %) of MFI or CV in Th1 per Th0 was calculated for each signaling pathway (Fig 10). A higher ratio of MFI (higher value of increase% of MFI) and lower ratio of CV (lower value of increase% of CV) was a more attractive signaling pathway, indicating that the right-side/lower position of the ball indicates an attractive signaling pathway (Fig 10). If the position of the ball is almost the same, the smaller size of the ball is a more attractive signaling pathway (lower fluctuation) than a larger ball.

**Fig 10.**
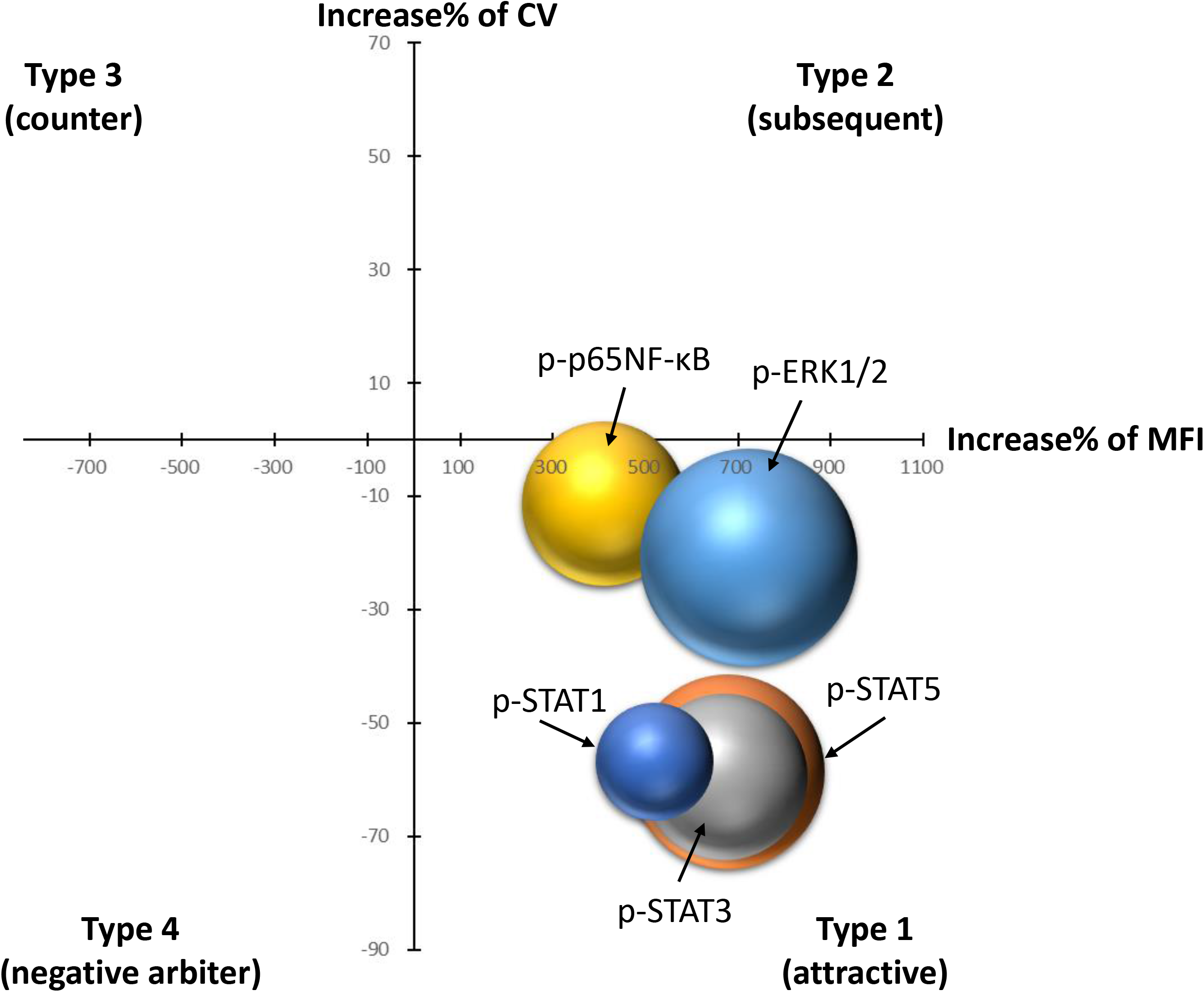
Comparison of contribution to T cell differentiation among signaling pathways using increase in the percentage (increase%) of MFI and CV. After the differentiation of T cells induced by OVA stimulation, MFI and CV of pSTAT1, pSTAT3, pSTAT5, pERK1/2, and p-p65NF-κB were measured as shown in Fig 9. The increase in the percentage of MFI and CV and the diameter of bubble size were calculated as follows: increase% = (Th1 – Th0)/Th0 × 100; 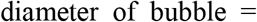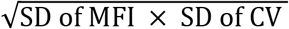, where SD is standard deviation. The horizontal axis represents the increase% of MFI, and the vertical axis is the increase% of CV. The bubble chart figure was drawn with Excel software (version 365).

As shown in Fig 10 and Fig EV5, pSTAT1, pSTAT3, and pSTAT5 were more attractive pathways for Th1 differentiation compared to ERK and NF-κB pathways. Furthermore, pSTAT1 was the smallest size of ball among pSTATs. Previously, the importance of STAT signaling pathways in T cell differentiation has been reported, and STAT1 contributes to initial Th1 commitment (Hibbert *et al*, 2003; Rochman *et al*, 2009; Takeda *et al*, 2003). Therefore, this analysis suggested that the categorization may be applied to detect the contribution to a hub signaling pathway, such as T cell differentiation.

## Discussion

Activation energy and fluctuating state in cell signaling were detected as MFI and CV by FCM. Changes in MFI and CV are utilized for a recognition of signal characters, such as “attractive,” “subsequent,” “passive,” “counter,” and “negative arbiter”, and these categorizations do not directly correspond to robust, regulated, oscillatory, and homeostatic signaling pathways, respectively, but are “elements” that form signaling pathways.

The robust signaling pathway is recognized as a relative decrease in CV; hence, it is recognized as the relationship between “passive” and “attractive (or negative arbiter)” The regulatory signal, which includes a relative increase in CV, is recognized by the relationship between “passive” and “subsequent (or counter)”. Oscillating signaling pathways are composed of subsequent, and intermediate between counter, and negative arbiters. The homeostatic signaling pathway is an oscillatory pathway between “passive” and “counter” and is, thus, detected as “passive” with a correlation between MFI and CV. Furthermore, during the observation period of cell activation in case of simple signal activation, MFI and CV showed a positive or negative correlation.

Many researchers may wonder what to consider when the histogram is bimodal or distorted. In this study, the shape of the histogram was topologically transformed to a simple shape depending on the change in MFI and CV, as shown in Table 1. The histogram was bimodal or distorted, suggesting that the cell population had several cell subsets depending on the response to the stimulants. A bimodal or distorted histogram will change the CV depending on the ratio of the cell subset in the cell cluster.

The existence of type 3 with a negative correlation between MFI and CV (homeostatic ERK pathway) allowed us to speculate that there is a mirror image of type 3 with a positive correlation between MFI and CV. The existence of an increase in CV with no change in MFI (p-tyrosine) also speculated that there is a mirror image of the existence of a decrease in CV without a change in MFI. Furthermore, the existence of a decrease in MFI with no change in CV (p-p65NF-κB in later phase) also induced the speculation of the existence of an increase in MFI without a change in CV. These patterns imaging using statistical changes in MFI, CV, and their correlations can logically make six variations of type 3 (Fig 11). The six variations of type 3 may also be elements for signaling pathways.

**Fig 11.**
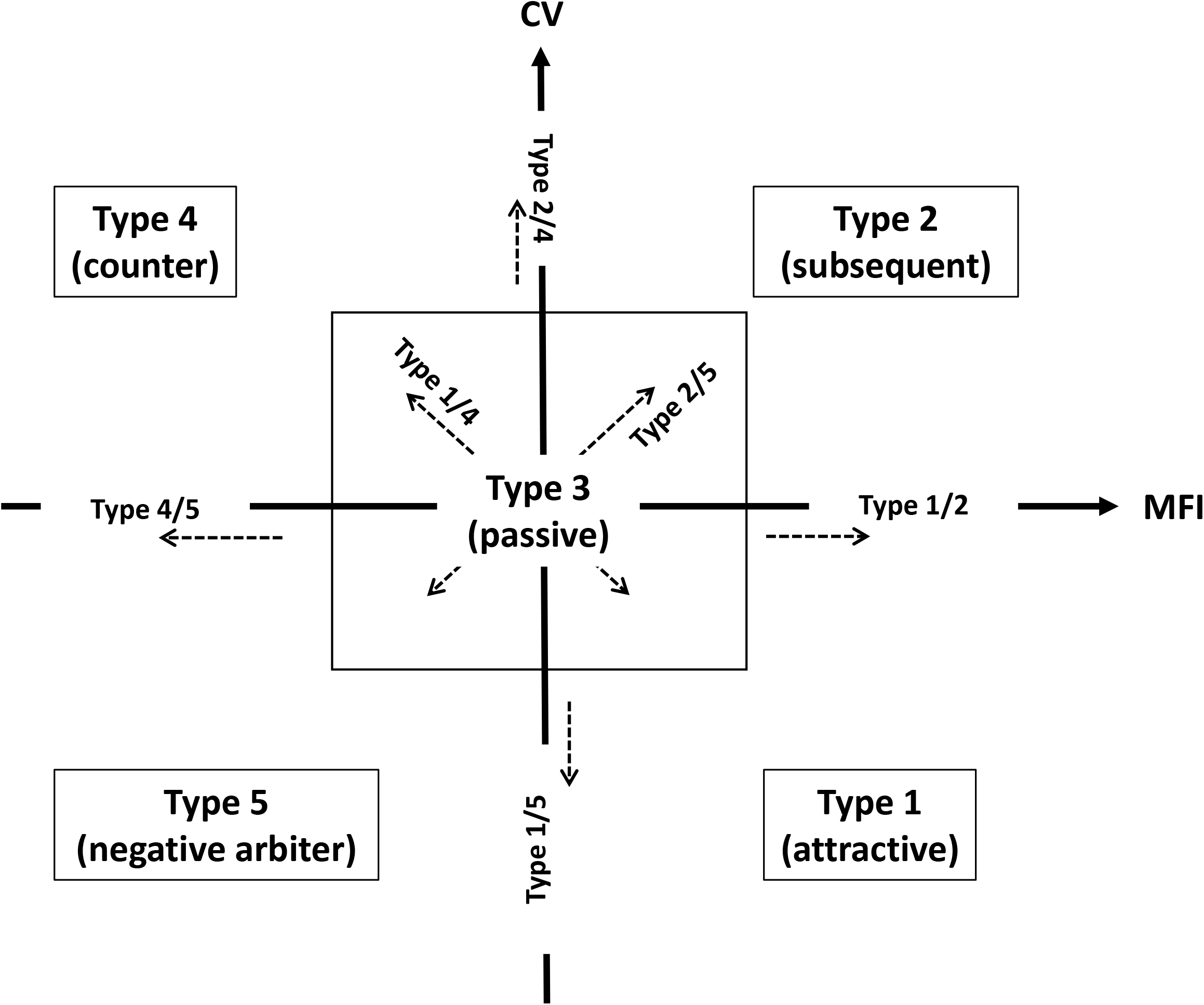
Schematic representation of six separations of type 3 (passive) with the five main categorizations. Moving energy and fluctuation are shown as two-dimensional axes, which are increases in MFI (horizontal axis) and CV (vertical axis), respectively. The five main categories are indicated in each rectangle. The six separations of type 3 are shown as dashed arrows. Eleven types of cell behavior may indicate elements of signaling pathways.

There are numerous combinations of signaling pathways, stimulants, and cells. Many researchers may feel that the number of examined patterns may not be sufficient to verify the categorization. However, when the phenomenon is recognized by activation or fluctuation, such as MFI or CV, there are only 11 patterns, logically. Furthermore, both activation and fluctuation are relatively determined by comparison. For example, the STAT1 signal is not always robust and is more robust than other signals. In other words, these are relative division but not absolute division. Therefore, we believe that this study, which shows the existence of 11 patterns, is necessary and sufficient.

According to the results of differentiated T cells in this study, the hub signaling pathway for differentiation was discriminated by “attractive” pattern. A simulation analysis with a focus on the “attractive” pattern may easily clarify the hierarchical structure of the ontology related to hub phenomena in trans-omics analysis. Discover an observable viewpoint to detect robustness means that we are able to define an emergence of phenomenon in cell population.

In this study, we proposed novel methods for recognizing signaling pathways using FCM. The contraction and fluctuation of cell clusters are useful for categorizing the signal characteristics. Previously, we demonstrated that CV of the neutrophil surface molecule GPI-80 was useful for the detection of neutrophil differentiation and heterogeneity (Kato *et al*, 2019; Takeda *et al*, 2016). These observations suggest that the categorization will be not only used for phosphorylated molecules but also cell surface molecules. We expect that this method can be applied to various molecule statuses and that it will evolve into an effective technique with various simulations for application in systems biology.

## Materials and Methods

### Human peripheral blood

This study was approved by the Ethics Committee of Yamagata University Faculty of Medicine (approval number: H28-265). Peripheral blood was collected from healthy volunteers after obtaining informed consent and heparinized with 5 U/mL of low molecular weight heparin. The average age of the volunteers was 47.0 ± 1.6 years (n = 5, male). Following sample collection, the blood was immediately transferred into 1.5-mL microtubes and used for experiments.

### Mice

OT-II transgenic mice (OT-II mice), which express T cell receptor (TCR) α and β chains that recognize the MHC class II I^b^-restricted OVA peptide (residues 323–339) in a C57BL/6J background, were kindly provided by Dr. W. Heath (WEHI, Melbourne, Australia). The mice were bred at the animal facilities of Yamagata University, Faculty of Medicine, under specific-pathogen-free conditions and were used for experiments at 6–12 weeks of age. The animal experiments were approved by the Animal Experiment Committee of Yamagata University, Faculty of Medicine (approval number, 31006).

### Cell lines

HL-60 cells were obtained from JCRB Cell Bank of National Institutes of Biomedical Innovation, Health and Nutrition. (Tokyo, Japan). U937 cells were gifted by Prof. N. Ishii (Department of Immunology, Tohoku University School of Medicine). These cells were maintained in RPMI 1640 (Sigma-Aldrich, St. Louis, MO, USA). The culture media were supplemented with 10% heat-inactivated FCS (Biowest, Nuaillé, France), 50 U/mL penicillin G potassium, and 50 μg/mL streptomycin sulfate. These cell lines were cultured at 37°C in 5% CO_2_ at high humidity.

### Antibodies

Antibodies used in this study were obtained as follows: Pacific Blue-conjugated anti-human CD3 mAb (UCHT1), PE/Cy7-conjugated anti-mouse CD4 mAb (GK1.5), PE-conjugated anti-T-bet mAb (4B10), PE-conjugated anti-Bcl-6 mAb (7D1), and Alexa647-conjugated anti-phosphorylated tyrosine mAb (PY20) from BioLegend (San Diego, CA, USA); FITC- or PE-conjugated anti-human CD14 mAb (MψP9), V450-conjugated anti-mouse CD3 mAb (17A2), PE-conjugated GATA3 mAb (L50-823), PE-conjugated anti-RORγt mAb (Q31-378), Alexa488-conjugated anti-pY701-STAT1 mAb (4a), Alexa488-conjugated anti-pY705-STAT3 mAb (4/P-STAT3), Alexa488-conjugated anti-pY694-STAT5 mAb (47/Stat5), Alexa647-conjugated anti-pT202/pY204-ERK1/2 mAb (20A), V450-conjugated anti-pS473-Akt mAb (M89-61), and Alexa488-conjugated anti-pT180/pY182-p38MAPK mAb (36/p38) from BD Biosciences (San Jose, CA, USA); PE-conjugated anti-GPI-80 mAb (3H9) from MBL (Nagoya, Japan); Alexa647-conjugated or non-conjugated anti-pS536-NF-κBp65 rabbit mAb (93H1) from Cell Signaling Technology (Beverly, MA, USA); and Alexa488-conjugated goat anti-rabbit IgG Ab from Thermo Fisher (Molecular probes, Eugene, OR, USA). For the mouse control mAbs, IgG1 (MOPC-21) and IgG2a (G155–178), were obtained from BD Biosciences and BioLegend.

### Cytokines and stimulants

IFN-α and G-CSF were gifted by Takeda Pharmaceutical Co. Ltd (Osaka, Japan) and Chugai Pharmaceutical (Tokyo, Japan), respectively. IL-21 was purchased from Peprotech (Rocky Hill, NJ, USA). LPS (from *Escherichia coli* O127:B8) and PMA were purchased from Sigma-Aldrich. OVA was obtained from FUJIFILM Wako Pure Chemical Corporation (Osaka, Japan).

### Stimulation of cell lines and whole blood cells

Cell stimulation was performed as previously reported (Takeda *et al*, 2014). Briefly, HL60 and U937 cells (2 × 10^5^ cells/mL, 1 mL/tube) were preincubated in microtubes for 60 min at 37°C in RPMI1640 containing 10% FCS. Whole peripheral blood cells were transferred into microtubes (0.3 mL/tube) and preincubated for 20 min at 37°C. After the addition of various stimulants, the microtubes were immediately vortexed for a few seconds and incubated at 37°C using a heat block. After the stimulation, HL60 and U937 cells (1 mL) were immediately fixed by addition of 0.2 mL of x5 Lyse/Fix buffer (BD Biosciences, San Jose, CA, USA) for 10 min at 37°C. Whole blood cells (0.1 mL) were also immediately fixed by suspension in 1.4 mL of x1 Lyse/Fix buffer for 10 min at 37°C. The fixed cells were packed by centrifugation at 800 ×*g* for 1 min at 24°C and then stocked in 90% methanol (0.3 mL/tube) at −20°C until staining with antibodies.

### T cell stimulations

Spleens from OT-II mice were minced and homogenized, and the cell suspensions were suspended in RPMI1640 medium containing L-glutamine and 25 mM HEPES and supplemented with 10% (v/v) fetal calf serum (FCS), 50 μM 2-mercaptoethanol, 100 units/mL of penicillin, and 100 μg/mL of streptomycin. The cells (4 × 10^6^ cells/mL) were incubated with OVA (0.2 mg/mL) for 7 days. After incubation, the cells (1 mL) were fixed with the addition of x5 Lyse/Fix buffer (0.2 mL) for 10 min at 37°C. The fixed cells were packed by centrifugation at 800 ×*g* for 1 min and then stocked in 90% methanol (0.3 mL/tube) at −20°C until staining with antibodies.

### Cell staining with various antibodies and FCM analysis

The fixed cells were washed with 0.8 mL phosphate-buffered saline (PBS) and suspended in PBS containing 3% FCS and 0.1% sodium azide. The cells were incubated with each antibody for 45 min at −22–25°C after FcR blocking using Fc blocker (Human TruStain FcX, Biolegend) or anti-mouse CD16/CD32 antibody, clone 2.4G2. After the reaction, the cells were washed with PBS and measured by flow cytometry (ec800, Sony, Tokyo, Japan; FACSCanto II, BD Biosciences, Franklin Lakes, NJ, USA). Cell debris were excluded from the analysis by forward- and side-scatter gating. MFI and robust CV were analyzed using the FlowJo software (version 7.6.5, TreeStar, Ashland, OR, USA). FlowJo software calculates the robust CV using the formula as follows: CV = 100 × 1/2 [(Intensity at 84.13 percentile) – (Intensity at 15.87 percentile)] / Median. This calculation is not affected by the number of cell events.

### Statistical analysis

Data are presented as mean ± standard error (SE). Statistical analysis was performed using the Prism Software (version 5.03, GraphPad Software, San Diego, CA, USA). *p* < 0.05 was considered statistically significant.

### Data availability

The source data for all figures are provided as Supplementary documents. The representative analysis using flow cytometry data are provided as Expanded View Figures. The raw flow cytometry data for all figures are available upon request.

## Acknowledgements

This work was supported by a Grant-in-Aid for Scientific Research No. 22590432. We would like to thank Associate Professor Yutaro Obara (Department Pharmacology, Yamagata University Faculty of Medicine) for valuable suggestions on ERK pathways.

## Author contributions

Y.T. conceived and initiated the study. Y.T., K.K., R.M., and H.A. designed the experiments. Y.T., K.K, R.M., and S.S. performed the experiments. Y.T. and K.K. analyzed the data. Y.T. and H.A. wrote the manuscript with input from all authors.

## Expanded View Figure legends

Fig EV1. Representative gating strategy of HL60 cell samples (A – C). HL60 cells were measured by flow cytometry (ec800). The panel of FS-Lin vs SS-Lin (A) shows the gate for exclusion of cell debris and aggregated cells. The gated cell population was expressed on the histograms of phosphorylated STAT3 (pSTAT3) data, which are shown as pSTAT3 (horizontal axis) in (B) or (C). The gray-filled histograms are initial state of the cells (0 min), and the open black histograms in (B) or (C) indicate the stimulated cells for 10 min with 300 U/mL IFN-α or 3 nM G-CSF, respectively.

Representative gating strategy of U937 cell samples (D – F). U937 cells were also measured by flow cytometry (ec800). The panel of FS-Lin vs SS-Lin (D) shows the gate for exclusion of cell debris and aggregated cells. The gated cell population was expressed on the histograms of phosphorylated p65NF-κB (p-p65NF-κB) data, which are shown as p-p65NF-κB (horizontal axis) in (E) or (F). The gray-filled histograms are initial state (0 min) of the cells, and the open black histograms in (E) or (F) indicate the stimulated cells with 100 ng/mL PMA for 30 min or 120 min, respectively.

The raw data of histograms (mean fluorescence intensity, MFI; and robust coefficient variation, CV) are presented in the rectangle at right side of each histogram panel.

Fig EV2. Representative gating strategy for T cells or neutrophils. Whole blood cell samples were measured by flow cytometry (FACSCanto II). The panel of FSC-H vs FSC-W (A) shows the gate for exclusion of erythrocytes and the ghost cells, and then the gated cells were re-gated by the panel of SSC-H vs SSC-W (B), which also indicates the gate for exclusion of cell debris and aggregated cells. The panel of CD3 vs GPI-80 (C) shows the gate for the selection of CD3^+^ T cells or GPI-80^+^ neutrophils. The T cell population or neutrophil population were expressed on the histograms of phosphorylated ERK1/2 (pERK1/2) data, which are shown as pERK1/2 (horizontal axis) in (D) or (E), respectively. The gray-filled histograms are initial state (vehicle) of pERK1/2, and the open black histogram are indicated as stimulated cells with 100 U/mL IFN-α + 1 nM IL-21 + 1 nM G-CSF for 30 min. The open black histograms are almost overlapped on the gray-filled histograms in both (D) and (E).

Fig EV3. Representative gating strategy for monocytes. Whole blood cell samples were measured by flow cytometry (FACSCanto II). For exclusion of erythrocytes, the ghost cells, cell debris and aggregated cells, gating were performed on FSC-H vs FSC-W and SSC-H vs SSC-W panels as same as Fig EV2. For the selection of monocytes, the panel of CD14 vs CD3 (A) or CD14 vs AmCyan Background staining (B) shows the gate for the selection of monocyte cells. The monocyte population was expressed on the histograms of phosphorylated ERK1/2 (pERK1/2) (C), phosphorylated tyrosine (pY) (D), phosphorylated Akt (pAkt) (E) or phosphorylated p38MAPK (p-p38MAPK) (F). The gray-filled histograms are the treated cells with vehicle [LPS (−)] for 30 min, and the open black histogram are indicated as the stimulated cells with LPS [LPS (+)] for 30 min. The raw data of each histograms (MFI and CV) are presented in the rectangle at right side of each histogram panel.

Fig EV4. Representative gating strategy for helper T cells (CD3^+^CD4^+^ T cells). Splenocyte samples were measured by flow cytometry (FACSCanto II) after stimulated with ovalbumin for 7 days. The panel of FSC-H vs FSC-W (A) and the panel of SSC-H vs SSC-W (B) are the gate for exclusion of cell debris and aggregated cells. The panel of FSC-A vs SSC-A (C) shows the gate for the selection of lymphocytes. The panel of APC-Cy7-background vs PerCP-Cy5.5-background (D) is the gate for the exclusion of cells which have strong intrinsic fluorescence. The CD3^+^CD4^+^ helper T cell population was selected as shown in (E), and then separated to T-bet^+^ Th1 cells and T-bet^−^ Th0 cells. These cell subsets were expressed on the histograms of phosphorylated STAT1 (pSTAT1) (G), phosphorylated STAT5 (pSTAT5) (H), or phosphorylated p65NF-κB (p-p65NF-κB) (I). The gray-filled histograms are Th0 cells, and the open black histogram are indicated as Th1 cells. The raw data of each histograms (MFI and CV) are presented in inside of each histogram panel.

Fig EV5. Analysis of STAT1, STAT5, and NF-κB signaling pathways in Th0 and Th1 cells using FCM. Splenocytes were stimulated with ovalbumin for 7 days. After the stimulations, the cells were fixed/permeabilized and stained with mAbs (V450-conjugated anti-mouse CD3 mAb, PE-Cy7-conjugated anti-mouse CD4 mAb, Alexa647-conjugated p-p65 NF-κB mAb, PE-conjugated anti-T-bet mAb, Alexa488-conjugated pSTAT1 mAb, or Alexa488-conjugated pSTAT5 mAb or a mixture of PE-conjugated anti-T-bet mAb, anti-GATA3 mAb, anti-RORγt mAb, and anti-Bcl-6 mAb). Cells were analyzed by FCM (FACSCanto II). Th0 cell subset was gated as T-bet^−^GATA3^−^RORγt^−^Bcl-6^−^ in CD3^+^CD4^+^ cells (open circle), and Th1 cell subset was gated as T-bet^+^ in CD3^+^CD4^+^ cells (closed circle). MFI and CV of pSTAT1 (A), pSTAT5 (C), and p-p65 NF-κB (D) are presented. Data were obtained from three independent experiments, and the statistical significance was calculated by paired Student’s *t*-test. \**p* < 0.05; ns, not significant.

